# Cloud-enabled Fiji for reproducible, integrated and modular image processing

**DOI:** 10.1101/2021.10.22.465513

**Authors:** Ling-Hong Hung, Evan Straw, Shishir Reddy, Robert Schmitz, Zachary Colburn, Ka Yee Yeung

## Abstract

Modern biomedical image analyses workflows contain multiple computational processing tasks giving rise to problems in reproducibility. In addition, image datasets can span both spatial and temporal dimensions, with additional channels for fluorescence and other data, resulting in datasets that are too large to be processed locally on a laptop. For omics analyses, software containers have been shown to enhance reproducibility, facilitate installation and provide access to scalable computational resources on the cloud. However, most image analyses contain steps that are graphical and interactive, features that are not supported by most omics execution engines. We present the containerized and cloud-enabled Biodepot-workflow-builder platform that supports graphics from software containers and has been extended for image analyses. We demonstrate the potential of our modular approach with multi-step workflows that incorporate the popular and open-source Fiji suite for image processing. One of our examples integrates fully interactive Fiji macros with Jupyter notebooks. Our second example illustrates how the complicated cloud setup of an computationally intensive process such as stitching 3D digital pathology datasets using BigStitcher can be automated and simplified. In both examples, users can leverage a form-based graphical interface to execute multi-step workflows with a single click, using the provided sample data and preset input parameters. Alternatively, users can interactively modify the image processing steps in the workflow, apply the workflows to their own data, change the input parameters and macros. By providing interactive graphics support to software containers, our modular platform supports reproducible image analysis workflows, simplified access to cloud resources for analysis of large datasets, and integrated workflows across imaging, genomics and transcriptomics data.

## 1 Introduction

The coupling of imaging data with localized clinical omics data in emerging fields such as spatial genomics is opening new frontiers in biomedicine. Computational techniques are essential to the extraction of biological knowledge from imaging data that are rapidly increasing in scope and scale [1]. Modern image datasets are digital, can span both spatial and temporal dimensions with additional channels for fluorescence and other data giving rise to datasets that can be terabytes in size. Image analysis suffers from many of the same problems plaguing other big data fields including scalability, reproducibility, interoperability, and accessibility. Analytical workflows for processing images are becoming increasingly complex requiring the orchestration of multiple pieces of software, which raises problems of automation and versioning. Similar reproducibility and scalability problems exist in the analysis of omics data which have been largely solved by workflow engines using software containers and cloud computing. These engines have not been widely applied to image analyses as they are designed for batch operations, whereas image tools are often interactive and graphical. Instead, automation and batch processing in image analyses are mostly dependent on macros and scripts that allow multiple commands to be recorded and repeated. Expensive, and difficult to maintain workstations are often used to analyze large image datasets. These workarounds are not tenable in the long term as workflows grow more complex and data sizes continue to increase.

Our solution is the Biodepot-workflow-builder (Bwb) with interactive graphics support which we have extended for image analyses. Bwb is a containerized and easy-to-use graphical workflow engine. Bwb uses software containers represented by graphical widgets to precisely define the version and operating environment for each module. Reproducibility and reliability of workflows are enhanced as software containers can be updated and modified independently, without affecting other modules. Software containers greatly expand the scope of analyses possible, as modules are no longer restricted to a pre-defined software suite. Most importantly, data and software from other domains such as omics and statistical packages can be combined with image analyses. Software containers also facilitate cloud deployment, allowing users to easily scale the analysis for large datasets using on-demand cloud resources. This eliminates the need for access to expensive, dedicated local servers, making it possible for anyone with access to the cloud to conduct the analyses of large datasets. In this manuscript, we present a demonstration of the advantages of this approach with Bwb workflows using the open-source Fiji suite.

Fiji [2] is a widely used open-source software suite that extends the basic image processing capabilities of ImageJ [3] with an expansive set of plugins. Both ImageJ and Fiji support macros, which can record and automate a series of commands. Fiji’s graphical interface and macro recording feature make Fiji an easy to use image processing tool for both experienced and inexperienced users. While these features have contributed to Fiji’s widespread use, they have some important limitations. 1) Fiji lacks systematic version control making it difficult to reproduce workflows because the success and effect of macros can depend on the version of the Fiji plugin ensemble. 2) Fiji is designed for interactive, desktop usage making it difficult to execute on the cloud. Without the scalable processing capabilities available on the cloud, Fiji needs powerful local servers to process large datasets. 3) Image analysis workflows can include statistical and visualization tasks that are not readily available in Fiji. State-of-the-art statistical packages and visualization tools can facilitate integration with non-image datasets as in the case of spatial genomics.

### 1.1 Our contributions

We extended the Biodepot-workflow-builder (Bwb) platform [4] to support interactive and reproducible analyses of imaging data. Fiji can be run in Bwb as a standalone application on a laptop or on the cloud. Bwb can also automate imaging workflows where graphics need to be output and allow users to interrupt execution to interactively adjust and tweak processing parameters. With Bwb, multiple Fiji macros from different sources can be reproducibly chained together as the required versions of Fiji and Fiji plugins are encapsulated within each modular widget. Even plugins that require Fiji graphics to execute can now be part of an automated and reproducible workflow. Finally, computationally demanding image processing, such as light sheet microscopy, often consists of gigabyte or even terabyte-sized datasets, exceeding the capabilities of a laptop. Bwb is browser-based, such that Fiji and all its plugins run on the cloud exactly as they would on a laptop. Users do not need an expensive local server for memory and computationally demanding operations such as stitching together large multidimensional image datasets, but instead can leverage the on-demand scalable resources available on the cloud.

We have also built a open-source custom version of Bwb to demonstrate its image analysis capabilities. This version makes the Fiji module is available in the toolDock and includes the image analysis workflows described in this manuscript. The user starts Bwb, on a laptop, local server, or on the cloud and then connects to it using a browser. To use Fiji as a standalone, an icon is dragged onto the desktop and double-clicked to enter any startup parameters such as scripts and macro files. The Fiji widget can also be connected with other Bwb widgets to form complicated analysis workflows which can be saved and shared. The custom version of Bwb includes widgets for RNA-seq, DNA-seq, Jupyter notebooks, and for executing scripts in a variety of languages. Other workflows from Bwb can be loaded and their widgets used in custom workflows. The user can apply these workflows on their own data and the results can be saved. The customized Bwb container image also contains two workflows with new Fiji macros in Bwb, that demonstrate the utility of the optionally-automated, modular, containerized approach for image analysis.

## 2 Results

We describe our effort extending the Biodepot-workflow-builder (Bwb) platform to support Fiji in Section 2.1. With this Fiji integration, users can reproducibly execute imaging libraries, plugins, pipelines, and macros in Fiji on the cloud. In addition, Fiji can be combined with other Bwb containerized modules (such as Jupyter notebooks, visualization and genomic analysis widgets) to form multi-step workflows. Bwb and its workflows are executed inside downloadable containers, thus only requiring that Docker be installed to function. To run Fiji as a standalone application, the user launches Bwb and connects to the Bwb server using a browser or a VNC client. A toolDeck of widgets becomes available on the right including a Fiji widget. The Fiji icon is dragged onto the Desktop canvas. Double clicking on the icon reveals the forms for entering parameters and starting and stopping the execution. Upon beginning execution Bwb will automatically download and installing the application and its dependencies. The user interacts with Fiji as they would if Fiji were launched locally on a laptop. The user can stitch together multiple modules to create larger, more complicated workflows that can be saved and reproducibly shared with collaborators.

We demonstrate the utility of our approach by analyzing focal adhesions and stitching three dimensional (3D) images in Sections 2.2 and 2.3. The first workflow in Section 2.2 performs a multi-step, multi-executable analysis of focal adhesions. It utilizes a download widget, a Fiji macro for segmentation, and Jupyter notebooks for data analysis and visualization. Each module (widget) is isolated within its own software container, enhancing reproducibility and re-usability by insulating it from changes to other modules and the operating environment. The second workflow in Section 2.3 demonstrates how graphics enabled containerization can simplify the deployment of a computationally demanding workflow to a virtual machine of arbitrary size on the cloud. This workflow downloads the dataset and directory structure, executes a modifiable Fiji module to execute the BigStitcher [5] plugin for Fiji. The workflow can be run on the cloud, while retaining the interactivity and extending the functionality of Fiji executed on a local computer. With these two workflows, we demonstrate how our work enables scalable and reproducible bioimage informatic analysis, and also facilitates the integration of imaging data analysis with additional data types available through Jupyter notebooks and other Bwb genomics analysis modules.

### 2.1 Supporting graphical applications such as Fiji with software containers

The major technical challenge to porting desktop based image analyses applications to the cloud is to support the same graphical interface and display on the cloud that one would see on a laptop or desktop. Bwb supports two methodologies for accomplishing this using software containers [6, 7]. We combine both methods in Bwb to allow the user to export graphics from a container that functions both on a local laptop and on a remote cloud server. A container exports X11 commands to the Bwb desktop environment which draws the graphics to a framebuffer. The framebuffer is exported using the virtual network computing (VNC) [8] protocol to a viewer that displays the screen and manages user interactions. An open-source VNC (noVNC) viewer allows users to use their browser to directly interact with the cloud workflow and avoid the need to install additional VNC software. While the the basic X11/VNC mechanism works well for many graphical applications in the original Bwb, we made some adjustments to support Fiji. We also made some adjustments to Bwb to facilitate the creation of customized containers such as the one we have provided to demonstrate the Fiji workflows. Details of these modifications are provided in Methods.

### 2.2 Analysis of focal adhesions

Epithelial cell motility is crucial to many biological processes, including development, wound healing, and metastasis. It is driven by cell-substrate interactions, which are mediated by focal adhesions, large multi-protein complexes situated on the basal surface of cells and most often near the leading edge [9]. They indirectly link the actin cytoskeleton and extracellular matrix proteins. This enables them to translate actin bundle contraction into traction forces that pull them across a substrate [10]. Although all focal adhesions are composed of a core set of proteins, including vinculin and paxillin, there are significant compositional differences between cell types with over 100 distinct proteins having been reported to associate with focal adhesions [11]. Moreover, the numbers, sizes, and shapes of focal adhesions differ between cell types and are influenced by substrate stiffness, extracellular matrix, and signaling molecules, with these metrics being associated with cell motility [12]. Calculating the numbers, sizes, and shapes of focal adhesions requires the segmentation of these microscopic structures in immunofluorescent images. Since dozens or more focal adhesions can be present in each cell, this is a very laborious process. There is therefore a significant, cross-discipline interest in methods to automate focal adhesion segmentation [13]. However, because of differences between cell types, immunostaining methods, and imaging platforms and settings, this is not an easily standardizable task. Flexible methods for modifying existing segmentation models or generating new ones are desperately needed.

Our Bwb workflow and Fiji macros implement segmentation of focal adhesions using the algorithm described by Horzum *et al.* [14]. A set of images of fluorescently labeled paxillin available from the Focal Adhesion Analysis Server [13] was first downloaded using the Download Files widget in Bwb, along with the LoG3D plugin for Fiji [15], which implements a Laplacian of Gaussians filter. We implemented an option in the Fiji widget to specify the directory containing the LoG3D plugin (see Section 4.2).

A containerized copy of Fiji was then called to execute a Fiji macro that triggers actions to perform the image processing steps as described by Horzum *et al.* [14] that isolate the focal adhesions in the image. The macro then used the built-in “Analyze Particles” plugin to compute and save the area (in square pixels) and centroid coordinates for each focal adhesion to a CSV file. An ellipse was also fitted to each focal adhesion and the lengths of the major and minor axes of the ellipse as well as the angle of the major axis with respect to the horizontal were saved. Since Fiji’s macro language is intended to execute graphical user interface commands, the graphics and interface were exported from the Fiji container so the user can observe the status of the segmentation and interactively adjust the image in real time; see Figure 2 (B).

**Fig. 1.**
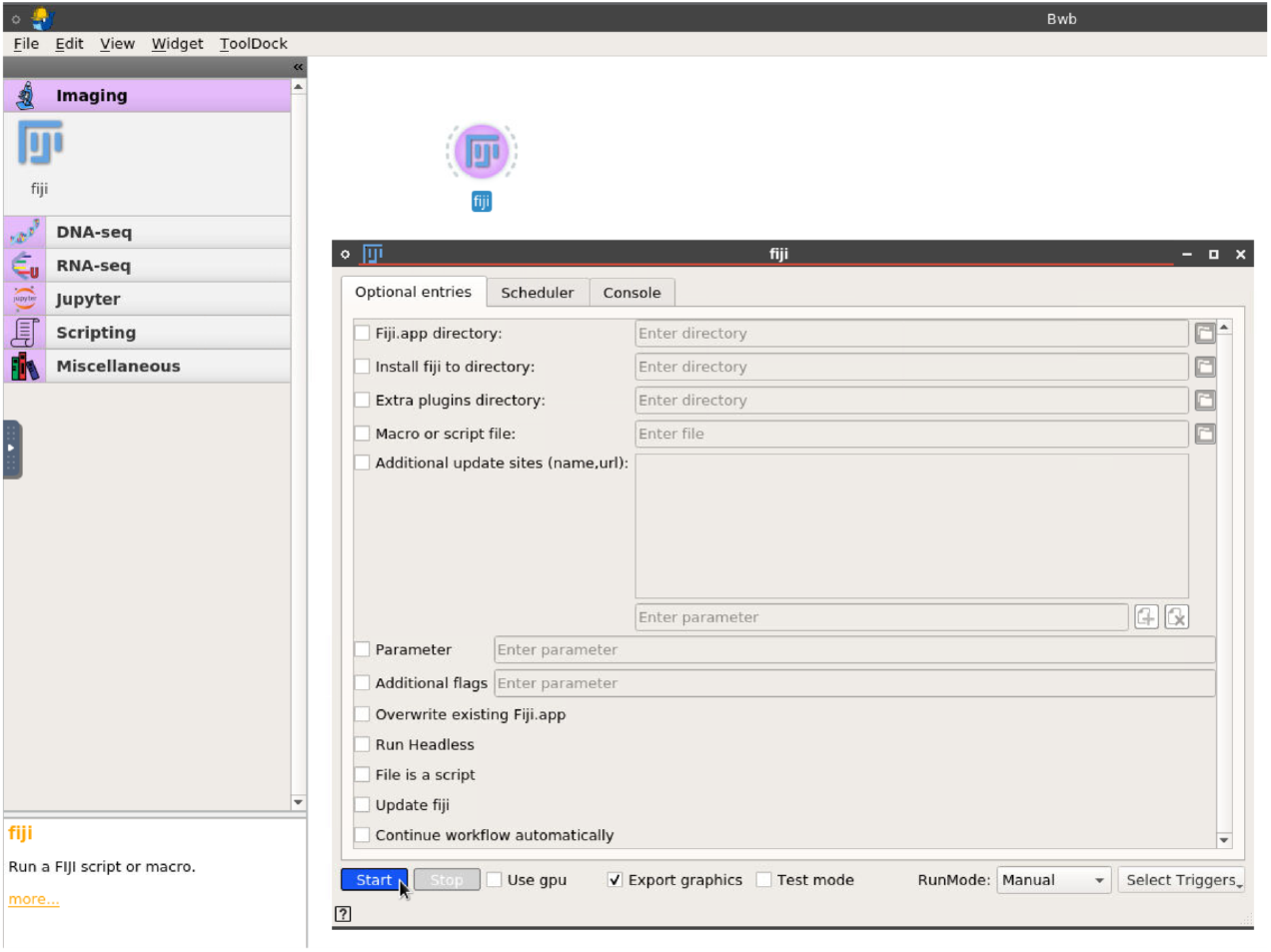
A screenshot of the custom version of the Biodepot-workflow-builder that demonstrates its image analysis capabilities. The Fiji icon (widget) has been dragged from the left sidebar Image Analysis drawer onto the canvas. The form-based interface for entering optional parameters to customize the Fiji suite is shown and is revealed by double-clicking on the dragged Fiji icon. The cursor is position over the start button which will launch the Fiji suite for standalone usage. Widgets from other drawers on the sidebar tool dock are revealed by clicking on the category and can be dragged to the canvas and connected together to form automated or semi-automated workflows for image analysis.

**Fig. 2.**
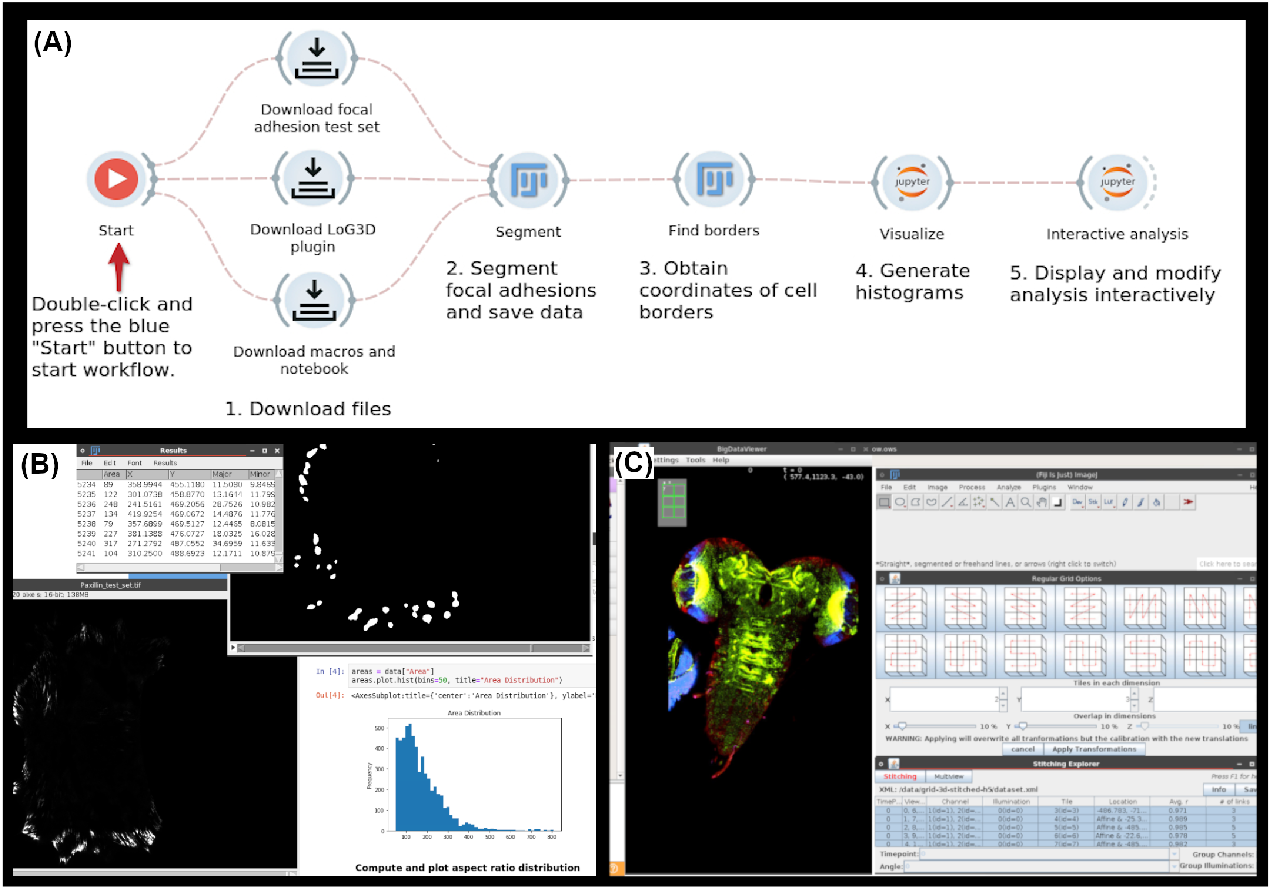
(A) The workflow for segmenting focal adhesions. Execution begins at the play button icon and proceeds to the right, with each widget being triggered upon completion of the task performed by the one to its left. (B) The workflow in (A) in action, with focal adhesion segmentation being performed in Fiji on the top and left, and results being displayed in Jupyter on the right. The top image window and table specifically show the results output by the “Analyze Particles” command; each white blob represents a particle, with statistics for each being shown in the table. (C) The left half of the image contains the stitched *Drosophila* nervous system displayed using BigDataViewer within the BigStitcher plugin. The right half of the image shows options for alignment to different regular grids as well as information for each tile separately.

Another macro was then executed in Fiji to identify the outline of the cell in each time-slice of the image; this is important to determine the orientation of each focal adhesion relative to the edge of the cell. The macro first performed automatic thresholding, and then performed a series of morphological operations to demarcate the cell. The operations consisted of morphological dilation, closure, filling holes using the Fiji “Fill Holes” plugin, and erosion to return the cell to approximately its original shape. It was determined interactively, that three rounds of the morphological operations were required to close holes and remove noise. The “Outline” operation in Fiji was then used to find the cell borders, which was saved as *xy*-coordinate pairs to a text file using the “Save XY Coordinates...” command.

Finally, a Jupyter notebook was executed to analyze and visualize the data from the previous steps. The Pandas [16] and NumPy [17] libraries were used to read the CSV file and obtain the segmentation and border coordinates. The Matplotlib [18] library was then used to create histograms to visualize the distribution of areas, aspect ratios (i.e. the ratio of the lengths of the major and minor axes), and the angles relative to the cell edge observed in the focal adhesions. Two separate widgets are used. The first widget loads all the required libraries and automatically executes the analysis to generate the final graphs. The second widget displays a copy of the executed notebook with the generated results. Either notebook can be altered which gives the user the ability to interactively explore different visualization and analysis options for a particular dataset and then decide whether to commit them to the automated part of the workflow to be applied to other datasets. See Figure 2 (A) for a screenshot of the complete workflow running in the Bwb platform and Figure 2 (B) for graphical output of this workflow. A video demonstration of this workflow is available at https://youtu.be/ymnmdRqS-pE.

### 2.3 Stitching of 3D images

Advances in microscopy techniques, such as lightsheet or confocal microscopy, have led to the generation of large overlapping three dimensional (3D) images. However, the raw data acquired with these microscopes are not directly suitable for visualization and analysis. Digital reconstruction of these 3D images require stitching and fusion of large numbers of overlapping image tiles. Additionally, these 3D datasets can be large, particularly for lightsheet microscopy which is often on the terabyte (TB) scale. This leads to computational challenges in terms of storage, efficient analysis, and visualization.

BigStitcher is a software package that enables interactive visualization, efficient image alignment and deconvolution of multi-tile and multi-angle image datasets [5]. Both alignment and viewing of terabyte-size datasets composed of overlapping three-dimensional (3D) image tiles are supported. BigStitcher offers options for fully automatic or interactive stitching. BigDataViewer [19] is provided with the package, for visualization of the aligned datasets. To accommodate the computational requirements, the original benchmarks for BigStitcher were performed using high-end local servers [5]. However, many researchers do not have access to this type of dedicated infrastructure but do have access to pay-as-you-go resources on the public cloud. We demonstrate how enabling execution of Fiji on the cloud democratizes access to computationally intensive applications such as BigStitcher.

BigStitcher is installed as a plugin through the Bwb Fiji widget. The demo workflow uses BigStitcher to align and display a multi-tile dataset of a 3D confocal scan of the nervous system of a *Drosophila* larva. The images consist of 6 tiles and 3 channels each. This data is available as an example dataset within BigStitcher’s Fiji plugin documentation and serves as a good example for the cloud capabilities of the software package. After opening the raw data in BigStitcher, the filename patterns for channels and tiles are selected by our macro. The images are then pulled up in the BigDataViewer plugin and aligned to a regular grid for easier viewing. Manual alignment is skipped in this workflow but is available as an option for more complex datasets that require user intervention in stitching. Figure 2 (C) shows the stitched image from the *Drosophila* dataset (123 MB) displayed in the BigDataViewer plugin. Stitching and viewing the raw data is completed in 20 seconds on a m5dn.4xlarge AWS EC2 instance. A video demonstration of this workflow is available at https://youtu.be/6S0KJEa3M0w.

## 3 Conclusions

Imaging analysis workflows often consist of disparate tools linked together in haphazard fashion and running on local hardware. This approach is characterized by two major challenges: 1) a lack of systematic version control, which compromises reproducibility, and 2) throughput that is limited by local hardware. To overcome these challenges we present an accessible, cloud-enabled and extensible platform. Notably, our platform also enables easy integration across multiple data types.

The Fiji image analysis suite has been containerized for reproducible deployment on local and cloud servers and in both cases functions in the same interactive manner as it does on a local host. Users can modify individual modules and corresponding input parameters in the workflow using a drag-and-drop user interface. Thus, the benefits of both interactivity and cloud capabilities are now accessible to non-programmers. We have demonstrated the utility of Bwb, a containerized, graphical cloud workflow execution platform for image analyses in two separate use cases. In these proof-of-concept use cases, we have connected Fiji to separate modules for downloading data, and integrated Fiji workflows with Jupyter notebooks for analysis and the visualization of results. In addition, we showed how our platform enables the execution of computationally expensive applications such as BigStitcher on the cloud.

The modularity, reproducibility, and accessibility of this approach democratizes both image analysis and the development of new analytical methods and packages. In addition, since our platform is open source and Bwb workflows can easily be shared, our platform enables researchers to easily reproduce the work of others or integrate new analyses into their own workflows. Moreover, the macros created in this work can be easily modified and adapted for different data sources and settings, making our approach highly extensible. Step-by-step descriptions of how this can be done have been published in our previous work [4]. Finally, Bwb also supports many genomics workflows [20] that can potentially be connected with image software like Fiji for interactive, cloud enabled spatial genomics analysis.

## 4 Methods

### 4.1 Implementation details for the customized Bwb container with Fiji

We also made two changes to Bwb to support the Fiji suite. In our original Bwb platform, some of Fiji’s menus were not displaying properly for some versions. This was overcome by modifying the VNC display settings. We also noted some problems that arose when multiple modules tried to draw to the same X11 screen. We modified Bwb so that each module would draw to its own X11 screen. These two changes were incorporated into the core Bwb platform allowing workflows incorporating the Fiji suite to be executed in the regular distribution of Bwb. In addition, we wanted to provide a customized Bwb version that had the Fiji widget available in the tool dock sidebar on startup and with the sample workflows available in the /workflows directory. We added some code to the core Bwb routines to allow the generation of customized Bwb containers with a small Dockerfile.

The actual construction of the Fiji widget was done using the standard mechanism for any Bwb widget. We first constructed a Fiji Docker container and uploaded it in our Biodepot repository for public downloads. The rest of the module is built using Bwb’s form based interface and can be easily changed and customized by the user if desired. This widget is saved as a directory of JSON and XML files. Bwb will automatically download or create the Fiji container if necessary and then Fiji will launch. Users can also load and mix and match different modules with Fiji to form new workflows using Bwb’s drag and drop interface, just as they can with Bwb widgets for statistical, genomics or transcriptomics analysis.

### 4.2 Macro for segmentation in the focal adhesion workflow

The focal adhesion segmentation workflow contains a Fiji macro widget which accepts a set of images (as a single TIFF file containing multiple pages) via a user parameter, and then performs the segmentation steps [14] on that set (see Section 2.2).

To allow for more interactive analysis, we have implemented a “continue workflow immediately” option in the Fiji widget. The default behavior is to exit Fiji immediately after execution and continue to the next step of the macro; however, when this option is deselected, Fiji will remain open after macro execution, allowing the user to analyze the results and/or edit the macros using all tools normally available to the user in Fiji. For example, at the end of the focal adhesion segmentation, the “Analyze Particles” command in Fiji is used, which detects distinct particles in an image and produces a black-and-white image where the particles are outlined and numbered, as well as a table containing the measured statistics for each numbered particle. By deselecting “continue workflow immediately”, the user could examine the results or the table, potentially to look for any noise mis-indentified as a particle (in which case the parameters of one of the intermediate steps of the segmentation will need to be changed). Additionally, the user could use Fiji’s built-in editor to edit the macro and make the desired changes to parameters. Upon exiting Fiji, the workflow will continue to the next step; the user may also choose to rerun the current step of the workflow with whatever new parameters they have defined.

Fiji macros can optionally accept a single string argument, which can be passed either from the command line or the runMacro command in Fiji, and can be retrieved within a macro using getArgument. The Fiji widget in Bwb has a parameter for passing this optional argument to a macro, which in the focal adhesion workflow is set to the file path of the image dataset to be segmented. Additionally, the workflow uses Bwb’s capability to pass values between widgets in a workflow to pass the extra argument to the border segmentation step as well, ensuring that the same dataset is used for each step, even if the file path is changed in the first widget. It is possible to change the dataset by first changing the URL in the downloader widget to the URL of a desired dataset, and then by changing the file path in the focal adhesion segmentation widget. This allows the workflow to be reused on different datasets with minimal changes to configuration.

### 4.3 Macro for stitching 3D images

The Bwb BigStitcher workflow contains a macro widget that accepts user parameters for defining an image dataset using BigStitcher. The dataset pattern must be specified in the widget, pointing BigStitcher to the channels and timepoints covered by the image dataset’s files. Additional parameters that can be defined include time-points per partition and the path to the input dataset for external testing. All the parameters within the BigStitcher demo workflow are automatically set on launch to work with the demo dataset. The macro widget uses these parameters to create a Fiji macro that will open the BigStitcher plugin, define a dataset according to the user parameters, and prepare the dataset for visualization using BigDataViewer within the BigStitcher plugin.

## Supplementary information

## Acknowledgments

This project has been funded in whole or in part with federal funds from the National Cancer Institute, National Institutes of Health, Department of Health and Human Services, under Contract No. 75N91020C00009 and 75N91021C00022. Hung, Reddy and Yeung are also supported by National Institutes of Health grant R01GM126019. We would like to thank Amazon Web Services for credits to Biodepot LLC for computing resources.

## Declarations

- Funding. LHH, SR and KYY are supported by NIH grant R01GM126019. LHH, ES, RS and ZC are supported by NCI SBIR contract 75N91021C00022.
- Conflict of interest/Competing interests. LHH and KYY also have equity interest in Biodepot LLC. KYY also received compensation from NCI SBIR contract numbers 75N91020C00009 and 75N91021C00022.
- Ethics approval. Not applicable.
- Consent to participate. Not applicable.
- Consent for publication. All authors have read and approved this article.
- Availability of data and materials.
- Code availability. Source code and documentation are publicly available at project web site https://github.com/BioDepot/fiji-demo.
- Authors’ contributions. This study was conceived and designed by L.H.H. and K.Y.Y. Z.C. designed and provided the background for the focal adhesion workflow and Fiji. E.S. implemented the first version of the Fiji widget for the focal adhesions workflow. S.R. implemented the BigStitcher workflow. E.S. and S.R. performed empirical experiments, created figures and videos in this manuscript. R.S. contributed to the implementation of graphics support in the Bwb. L.H.H. improved and tested all implementations in this project. K.Y.Y. drafted the manuscript. All authors contributed to the writing of the manuscript.

